# POLARIS: path of least action analysis on energy landscapes

**DOI:** 10.1101/633628

**Authors:** Evan Seitz, Joachim Frank

## Abstract

Free-energy landscapes are a powerful tool for analyzing dynamical processes - capable of providing a complete mapping of a system’s configurations in state space while articulating its energetics topologically in the form of sprawling hills and valleys. Within this mapping, the path of least action can be derived - representing the most probable sequence of transitions taken between any two states in the landscape. In this article, POLARIS (Path of Least Action Recursive Survey) is presented as a dynamic, global approach that efficiently automates the discovery of the least action path on 2D energy landscapes. Important built-in features of this program include plotting of landscape trajectories and transition state theory diagrams, generation of text files with least action coordinates and respective energies, and bifurcation analysis tools that provide downstream versatility for comparing most probable paths and reaction rates.

## INTRODUCTION

The construction and exploration of energy landscapes is essential for understanding the complex dynamics of molecular systems (1). Since the initial underpinnings of potential energy surfaces over a century ago (2), many attempts have been made to articulate its required properties. Fundamentally, the free-energy landscape is understood to be an intrinsic property of a given molecular system, independent of the experimental method used to obtain it (3). As these systems dynamically operate between distinct conformational states over many time and length scales, any mapping to the corresponding energy landscape must also be multidimensional and hierarchical in nature (4). Within this landscape, the sprawling layout of energy hills and valleys governs the system’s navigational probabilities - with deep wells representing distinct conformational states, and peaks acting to constrain the transitions between them. Further, since specific sequences of conformations give rise to biomolecular function, functional dynamics should be accounted for within this descriptive topology (5).

For decades, a variety of methods have existed to analyze energy landscapes (6); including global optimization algorithms (7), transition state search schemes (8, 9), and co-ordinate transformations (10), to name a few. During that time, many landscapes were successfully modeled for protein folding and related dynamics (11–13), while applications to larger, more complex macromolecular assemblies remained in their infancy (4). In recent years, however, technological advances in single-particle cryo-EM (14–17) have opened the door for constructing the free-energy landscape across a number of these previously incomprehensible systems (18–21).

Cryo-EM allows macromolecules to be experimentally visualized en masse via electron microscopy after being rapidly frozen in vitreous ice (quenching) at a rate assumed to be faster than the overall reconfiguration time of the system. As a result, the ensemble of frozen molecules closely approximates the Boltzmann distribution of states of the system immediately prior to freezing (3). Virtually all projection directions of the particle are obtained across each conformation in its state space. These data are then analyzed by a new computational technique employing Manifold Embedding (19–21) to construct a lower-dimensional state space of the system from its experimental sightings. The number of particles found in each state is then compared to the occupancy of the most populated conformation, with this ratio converted into standard free-energy differences via the Boltzmann factor to create the free-energy landscape (18). In this way, the landscape’s minimum energy wells represent the system’s most highly-preferred conformations as witnessed during the experiment. Through this method, the potential now exists for creating the conformational free-energy landscape for any molecular machine. With this powerful tool, new methods must be devised to extract valuable information needed for further biological elucidation.

Existing on a scale where thermal and deterministic forces act on equal footing, these macromolecules are constantly buffeted by the random motions of nearby solvent molecules as they compete to internally perform the sequence of motions required to operate. As a result, transitions between distinct states occur via a series of thermally-driven steps. This diffusive process can occur through a multitude of structural pathways in the energy landscape, ultimately creating an ensemble of transitory routes between any two configurations (5). Each of these available pathways represent a unique sequence of conformational events - with the probability for a transition to take one such pathway dependent on both the surrounding highest-energy peaks (4) and the total integrated energy along that path’s diverse range of intermediate states.

Historically, such notions stem from Maupertuis’ *principle of least action* (22), which defined the natural tendency of a system to choose the path among all available paths connecting two specified points that ultimately minimizes its *action*. Maupertuis worked closely with Euler, who continued to advance this concept as a minimization of *effort* (23) - an expression corresponding to our present-day understanding of potential energy. From this, the *path of least action* can be defined and computationally generated for the energy landscape - representing the most probable sequence of unique conformational changes taken by the system to navigate between two distinct configurations.

Many general techniques already exist to algorithmically solve pathfinding problems - seeking to identify the best path between any two points as defined by some overhead metric (e.g., shortest or fastest route). In the majority of these, graph theory plays a prominent role, with hallmark algorithm’s such as Dijkstra’s (24) (and many variants (25– 27)) proven successful at solving these problems. However, these methods rely on the underlying assumption that the kernel being used (representing the distance between any two points) is deterministically correct. By calculating this kernel across every combination of points, a weighted graph is produced, where a value (weight) is given to every edge connecting the set of vertices in the graph. A collection of decisions are subsequently made to ultimately find the collection of intermediate vertices such that the sum of their constituent edge weights is minimized.

However, in situations where the value for each edge weight is not conveniently predefined (as is the case in energy landscapes), uncertainty arises from the theoretically infinite number of kernel functions one could use to define such a weighting. For example, Graph Spectral Image Processing (28) aims to construct an underlying graph connecting pixels with weights via a bilaterally-filtered Gaussian kernel incorporating pixel intensity and distance, while also requiring two additional parameters. Alternatively, edge weights can also be defined based on local pixel patches or overhead features (to name a few) (29). Ultimately, the appropriate definition of these edge weights is application-dependent (28), with the solution to the pathfinding problem changing as the underlying relationships between each set of points is altered. As such, using edge weights to find the absolute path of least energy (in the context of energy landscape problems) requires that the optimized output from one arbitrary weighting function be compared to an infinite number of competing weighting functions - presenting an endless search above the procedures underlying each graph-based approach.

Further, the very choice of overhead metric within the contextual demands of each underlying system is a source of uncertainty. In the case of the macromolecular assembly driven by thermal motion, such a metric should reflect the total integrated energy of each possible path, while also analyzing these paths within the landscape’s hierarchy of distinguished scales (macroscopic, mesoscopic, and microscopic (4)). Regardless, algorithms do exist that use broad, graph-based approaches to navigate between states in the energy landscape. One such tool is the Dijksta-inspired MEPSA (Minimum Energy Pathway Analysis (30)) which will be analyzed and compared in detail with our method.

POLARIS (Path of Least Action Recursive Survey) provides an alternative approach to the minimum energy pathfinding problem by avoiding the arbitrary assignment of edge weights and extending its methods outside the realm of graph theory. Instead, POLARIS aims to prioritize and isolate the most energetically favorable coordinates - as these represent highly occupied transit regions in the landscape on the macroscopic scale. This initial method reflects the paradigm used by hierarchical pathfinders (31), whereby the high-level overview route is assigned first by analyzing the most favorable transit locations in between. Between these anchors in the landscape, all permutations of a set of higher-order lines are drawn with the net energies compared from each. The algorithm then takes the resulting minimum energy discoveries as new inputs to itself to repeat this procedure recursively - breaking down each best global line approximation into continuously finer, mesoscopic subsections until the finest-granular, microscopic path is resolved.

## METHODS

POLARIS is a cross-platform, open-source program written with Python 3.x (32), a graphic user interface built using PyQt5 (33), plots drawn using Matplotlib (34), and operations performed with the aid of NumPy (35), Python’s Itertools, and Bressenham’s Line Algorithm (36). POLARIS has been designed as a user-friendly tool able to analyze transition pathways through 2D energy landscapes in pursuit of expediting the discovery of their most significant pathways.

By defining spatial subdivisions for isolating minima within the energy landscape (1. Image Segmentation) and creating the set of all possible permutations between them over a range of increasing line orders (2. Permutational Analysis), POLARIS is able to compare global path approximations, isolate favorable minima, and construct local solutions between them as it methodically implements its optimized outputs as inputs into itself (3. Branching Recursions). These steps are performed consecutively, and will be discussed in this order in the subsections below.

As a note, to ensure that this procedure can be performed regardless of data dimensions, the width and height of the original data file are first trimmed to the nearest even pixel when necessary. Next, maximum energy borders are added to the file; expanding one power of 2 above the landscape’s dimensions (e.g. for a 70 × 70 dimension landscape, the data is given a border of 29 pixels on each side, such that a 2^7^ × 2^7^ landscape is created, with the extra space filled in uniformly with the highest energy value obtained from the given file). All borders are removed at the end of the computation.

### 1 Image Segmentation

Image segmentation is the process of dividing the landscape into a number of equally-sized squares and recording the coordinates of the local minimum-energy values within each (Figure 1). The ‘segmentation depth’, *n*, defines the number of these subdivisions created, and thus, the number of local minimum-energy coordinates recorded. The user defines the set of segmentation depths, {*n*_*i*_}, to use within the parameters tab; e.g., {*n*_7_, *n*_5_} or {*n*_7_, *n*_5_, *n*_4_, *n*_3_}, et cetera.

**Figure 1:**
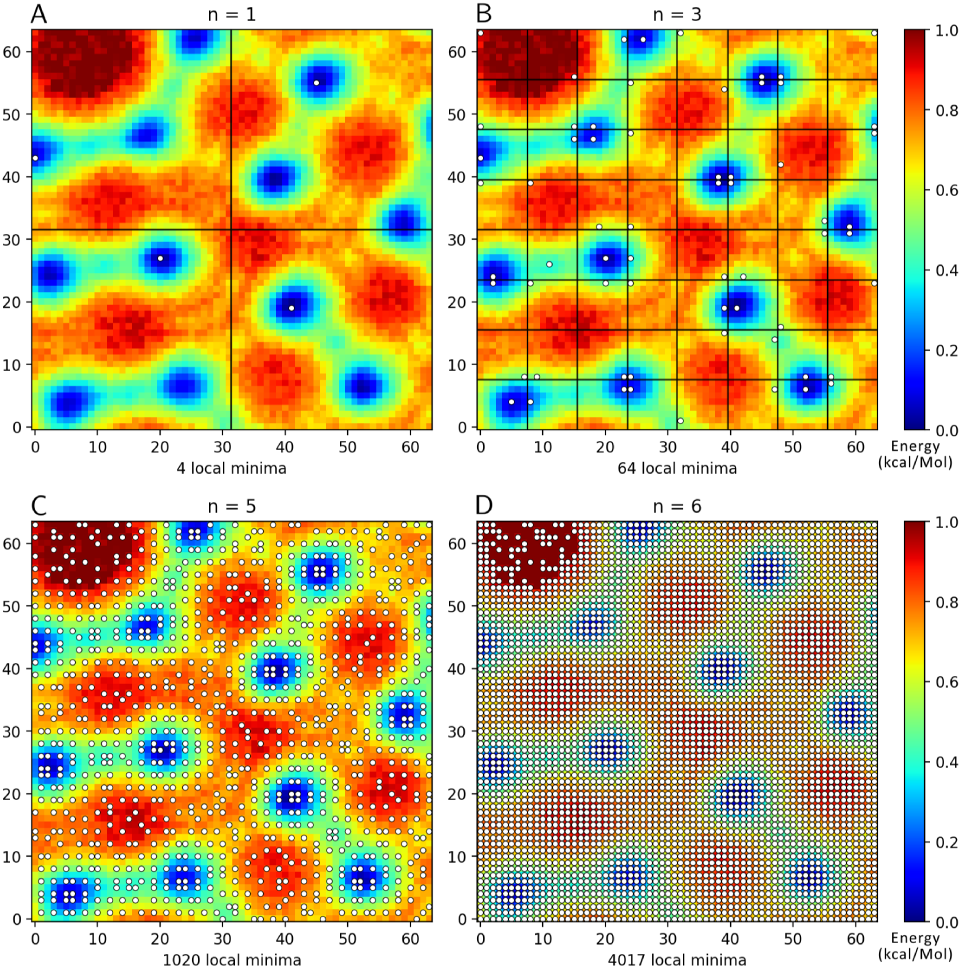
Computationally-generated landscape featuring increasing image segmentation depths, with segmenting lines (black) overlaid on *n*_1_ (A) and *n*_3_ (B) for division visualization. Local energy-minima are plotted as single white points exclusively occupying each square subdivision (encircled and numbered for clarity in A). In rare cases of multiple equal-valued minima per segment, only one minimum is selected, with subsequent depths capturing the alternative values. To maximize efficiency, no minima are obtained for segmented regions containing a uniform spread of globally maximum energy levels (as seen in the northwest corner of D). This landscape was created to exemplify the hierarchal scale seen in energy landscapes, whereby additive Gaussian noise was applied across all pixel intensities to represent unavoidable experimental uncertainties during data acquisition and processing.

For each 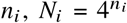 image subdivisions are created, with an equivalent number of minimum-energy points stored - such that, for *n*_1_, the coordinates of 4 local energy-minima are stored (Figure 1A). Within this set, the highest allowable value of *n*_*i*_, *n*_*max*_, is defined via that subdivision where further image divisions are impossible (e.g., for a 64 × 64 landscape, *n*_*max*_ = log_2_(64) = 6 subdivisions), whereby every segmented grid spans an area of exactly 1 pixel (Figure 1D).

As the segmentation depth is increased from *n*_1_ to *n*_*max*_, previously overlooked higher-energy minima (relative to the system’s global minima) are geometrically separated from the global minimum values and placed into their own neighboring grids. Since these locations become newly defined local energy-minima at higher depths, this method ensures that all points are eventually considered as potential nodes in the pathway permutational analysis to follow.

### 2 Permutational Analysis

Each set of 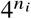 minimum-energy coordinates obtained from each *n*_*i*_ during image segmentation are subsequently used as transit options for comparing potential paths across the energy landscape. Here, for each *n*_*i*_, permutations of straight lines are connected between the set of 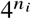 coordinates and the user-defined start point, *S*, and end point, *E* (Figure 2). The‘permutational order’, *r*, defines how many intermediate transit options can exist between *S* and *E*. For example, if there are 4 transit options (via *n*_1_), *r*_1_ would sample all ways of bridging *S* and *E* with straight lines using only 1 transit coordinate - a total of 4 permutations possible. These permutations are created via Itertools, such that for *n*_1_ with minima *M*_1_, *M*_2_, *M*_3_ and *M*_4_, paths *S* ↦ *M*_1_ ↦ *E, S* ↦ *M*_2_ ↦ *E, S* ↦ *M*_3_ ↦ *E* and*S* ↦ *M*_4_ ↦ *E* are generated (Figure 2A, Figure 2B).

**Figure 2:**
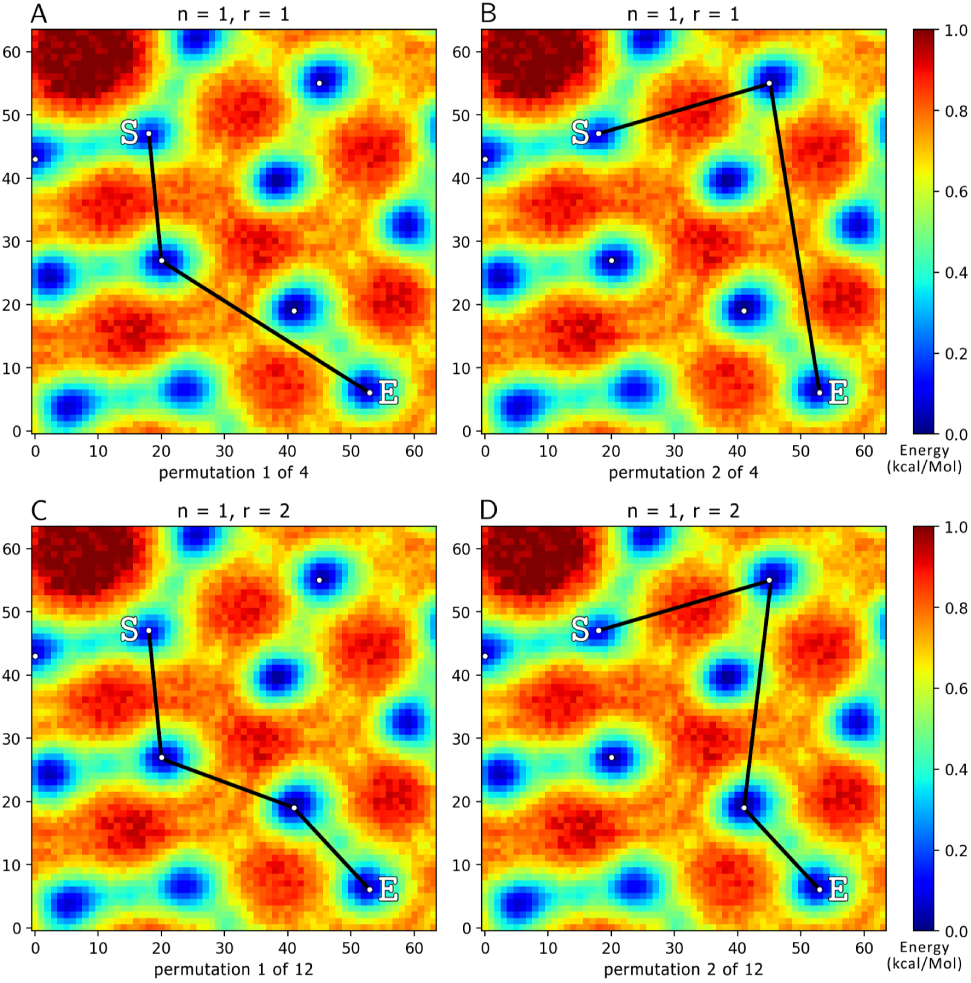
Computationally-generated landscape showing permutational order *r*_1_ (A, B) and *r*_2_ (C, D), each assigned *n*_1_. User-defined start and end points are labelled *S* and *E*, respectively. For *r*_1_, 4 permutations exist with one unique midpoint each, for which two have been shown (A, B). For *r*_2_, 12 permutations exist with two unique midpoints, for which two have been shown (C, D). The energies along the full set of these pathway permutations are integrated and compared (e.g., selecting the lowest-energy path of the 4 (A, B), or the lowest-energy path of the 12 (C, D), et cetera). The lowest-energy paths obtained from each {*n*_*i*_, *r* _*j*_}combination are then compared to find the lowest-energy approximation for all given combinations of permutational orders and segmentation depths.

For each user-defined value of *n*_*i*_, a partnering value for the permutational order, *r* _*j*_, must also be chosen; e.g., {(*n*_7_, *r*_1_), (*n*_5_, *r*_2_)}, et cetera. Here, *r* _*j*_, can range from *r*_0_ to *r*_*max*_, with *r*_0_ representing the line directly connecting *S* to *E* (with no midpoints in between). The value of *r*_*max*_ is restricted within the user interface to 5, with higher values deemed counterproductive to the aim of this interpolative procedure. Within this formulation, the standard permutation notation 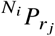 represents a unique combination of some *n*_*i*_ and *r* _*j*_, with 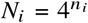. As an example, for *n*_2_ with *r*_2_, one such path *S* ↦ *M*_1_ ↦ *M*_16_ ↦ *E* is created amongst 239 other permutations, via *N*_2_ = (*n*_2_)^4^ = 16 and 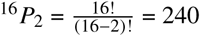.

Within each member of a given {*n* _*i*_, *r* _*j*_} permutational pool (e.g., 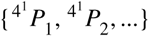, as seen in Figure 2), each set of ordered transit points are connected by straight lines; drawn using a variant of Bressenham’s Line Algorithm modified for energy awareness. These path approximations act to gather overhead awareness across different regions in the energy landscape. The total energies across the paths formed by each of these permutations is then independently integrated, with only the transit points from the minimum-energy permutation stored for that {*n*_*i*_, *r* _*j*_}. This process is then repeated for all combinations of {*n*_*i*_, *r* _*j*_}, ultimately only storing the set of transit points belonging to the lowest-energy path approximation discovered between the initial *S* and *E*. For example, if parameters {(*n*_7_, *r*_1_), (*n*_5_, *r*_2_)} are chosen, 16,384 + 1,047,552 = 1,063,936 paths are generated - with only the original list of minimum-energy coordinates pertaining to the path of overall lowest-energy stored. These minimum-energy coordinates are then introduced as inputs for the proceeding computations.

### 3 Branching Recursion

POLARIS next uses the discovered minimum-energy nodes from the previously performed permutational analysis recursively as inputs to itself (replacing the initial user-defined inputs *S* and *E* with the set of these newly discovered intermediate, minimum-energy nodes). First, a for-loop is created between each pair of intermediate nodes, such that if the pathway containing points *S* ↦ *M*_1_ ↦ *M*_2_ ↦ *E* were found as a minimum amongst all other permutations in the previously described steps, a loop containing *S* ↦ *M*_1_, *M*_1_ ↦ *M*_2_ and *M*_2_ ↦ *E* would emerge.

Within this loop, POLARIS performs branching recursion (Figure 3), repeating all of these aforementioned procedures on its newly obtained outputs. From here, the permutational steps are repeated for each new set of discovered start and end points - proceeding recursively down each inner branch until two output points having lowest line approximation *r*_0_ are found within each one of its individual leaves (defining the maximum extend of each branch). Upon each leaf break, the coordinate of that leaf is globally recorded. This process continues with the algorithm navigating throughout its recursive hierarchy until the path of least action is filled in completely with a set of coordinates spanning from the initial, user-defined start point to the initial, user-defined end point.

**Figure 3:**
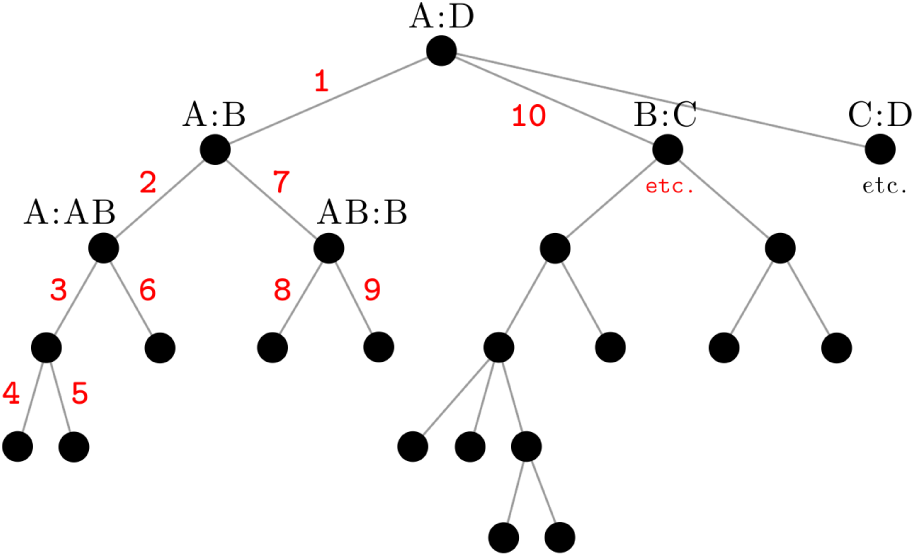
Schematic illustrating an example branching recursion hierarchy, given by initial seed A ↦ *D*. If the minimum-line outputs from the initial seed, `POLARIS(A,D)`, include points *B, C*, and *D*, `POLARIS(A,B)`, `POLARIS(B,C)`, and `POLARIS(C,D)` are subsequently called - with each one forming a new branch across the next-highest tier in the hierarchy. Red numbers indicate the order in which POLARIS navigates its recursive hierarchy. Broken leaves represent pairs of data points where further segmentation is futile; i.e., no lower energy line exists between them (occurring exclusively when *r*_0_ is returned as the lowest-energy approximation). As a note, the notation `POLARIS`(`input`1, `input`2) has been used here to describe one complete cycle of methods (1) and (2).

This process is ultimately akin to performing interpolation on the space between each start and end point within each recursion, whereby two anchors are known, with all points in between unknown. Those unknown points are then iteratively discovered via a complete set of the previously described permutational comparisons. Ultimately, these recursive steps break down the best global line approximation into continuously finer subsections within each recursion until a finest-granular path is resolved - giving rise to both global and local awareness of the landscape.

To remove unnecessary permutational calculations within the branching recursion hierarchy, and thus improve timing, once a set of {*n*_*i*_, *r* _*j*_} parameters are chosen, only those combinations within the set that are spatially viable for the current *S* and *E* coordinates are computed. As such, if the Euclidean distance between one set of *S* and *E* coordinates in the branch is smaller than the distance covered by the minimal spanning line of a specifically chosen combination of *n* and *r*, the permutational analysis for that *n* and *r* combination will be skipped.

### 4 Pathway Pruning

As an artifact of Bresenham’s Line Algorithm for drawing straight line approximations, there remain a multitude of ways to alias a diagonal line between the same two sets of coordinates (e.g., two different permutations can emerge for the same straight line when drawn *S* → *E* versus *E* →*S*). To work around this problem, pruning techniques have been implemented on the final set of coordinates of the complete least action trajectory. These include finite, local perturbations along each coordinate in the completed path. So long as these perturbations preserve the continuity of the overall path (no breaks), every point is iteratively sent to occupy its set of unrestricted neighboring pixels and reshuffle into those coordinates that ultimately minimize the pathway’s global energy.

As a note, any coordinates having zero-energy are never pruned from the pathway; safeguarded during this process for their significance in the energy landscape. This ensures that the overall line may still pass through these locations - including situations where the zero-energy point is neighbored by three other points in the path. Within this exception, it follows that no energy contribution is added to the path of least action.

## RESULTS

An experimentally generated free-energy landscape was used for testing POLARIS’ methods (Figure 4) - constructed through application of Manifold Embedding techniques to a set of ribosomal cryo-EM data (19). The ribosome has long been described as a “thermal ratchet machine” (37), whereby random energetic perturbations lead to large-scale shifts across its available configurations. The landscape obtained represents the conformational state space available to the naked ribosome (i.e., the ribosome in the absence of its functional ligands), with the reaction coordinates corresponding to the two highest-ranking, orthogonal factors for motion. These discovered coordinates capture the ribosome’s leading degrees of freedom during translocation - encompassing a composite of large-scale movements across its two subunits (19, 38), including rotations and swivels. Analysis of 3D reconstructions along the naked ribosome’s trajectory revealed that it undergoes conformational changes akin to those observed for factor- and GTP hydrolysis-driven translating ribosomes - clearly demonstrating the macromolecule’s intrinsically dynamic nature (3).

**Figure 4:**
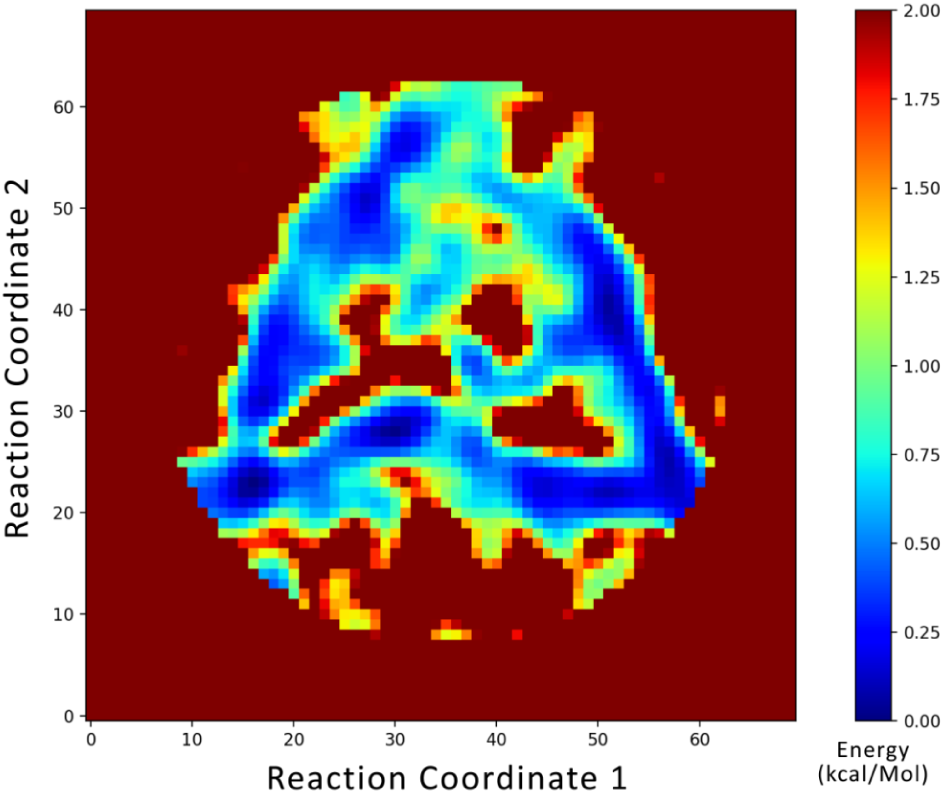
Energy landscape for the conformational coordinates of the naked ribosome during translation elongation. Data taken from ribosomal reaction coordinates (19).

Conformations reported by cryo-EM reveal that each “state” of the ribosome is actually an ensemble of structurally similar configurations clustered within a specific minimum-region on the energy landscape (3). Meanwhile, the rugged, higher-energy hills correspond to the less-favorable states of the macromolecular complex as it travels from one minima to the next. Given such a system, to properly delineate the ribosome’s key mechanisms of translation control, POLARIS was used to determine the most likely transitional pathways between different combinations of these free-energy basins.

### MEPSA Comparison

For validation, POLARIS’ results were compared to those obtained with a published energy landscape analysis algorithm entitled MEPSA (Minimum Energy Pathway Analysis) (30), which uses an approach similar to Dijkstra’s algorithm (24), with small differences in the sampling and traceback. The metric used to compare the two algorithms is the final integrated energy from a single continuous list of coordinates spanning between any two user-defined points.

The MEPSA tool allows two options for algorithmic comparison: the self-defined, less accurate ‘GLOBAL’ option (capable of accepting arbitrary user start and end inputs as well as predefined anchor points) and the more accurate option ‘NODE BY NODE’ (accepting only predefined global minima as user inputs). For the first comparison, the ‘GLOBAL’ approach was chosen to compare each program’s output given any set of user-defined start and end coordinates, here using (41, 51) and (17, 14) respectively on the ribosome energy landscape (Figure 4). Chosen for the complexity of the regions in between, this trajectory includes opportunities for traversing many narrow, branching low-energy pathways throughout the center of the landscape, as well as a final leap across a mountain range of highest-energy coordinates into a pool of minimum energy.

The net energy of the least-energy pathway found by MEPSA was 34.6 kcal/mol with a length of 106 points, containing many unnecessary regions where the algorithm appeared to draw jagged lines between its predefined nodes (Figure 5A). For the same arbitrary points, POLARIS returned a 78-point 24.7 kcal/mol path (Figure 5B), showing a difference of 9.9 kcal/mol between the two algorithms in favor of POLARIS.

**Figure 5:**
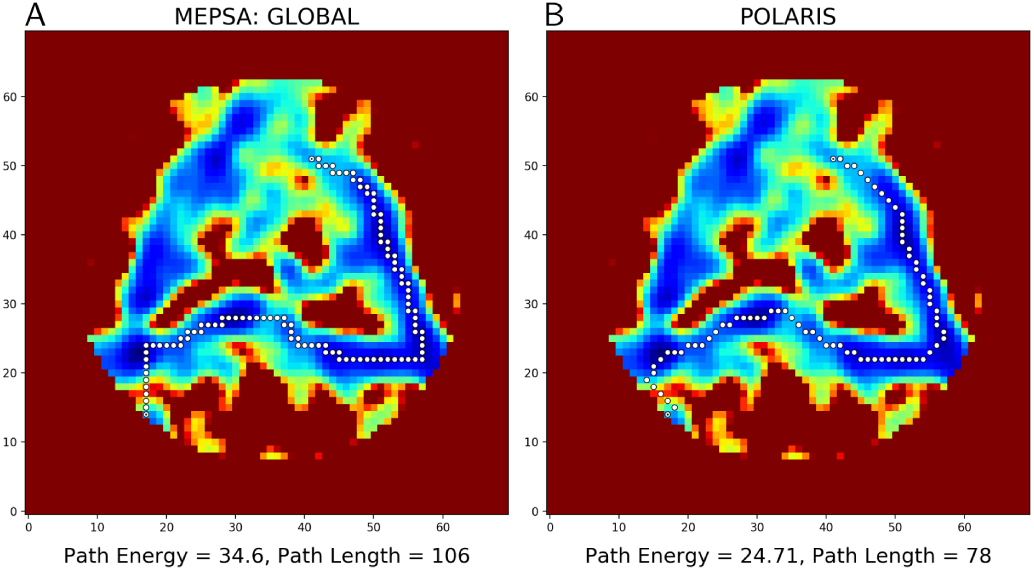
Comparison between MEPSA’s ‘GLOBAL’ algorithm (A) and POLARIS (B). While both algorithms found the same approximate global path, POLARIS returned a path of integrated energy 9.9 kcal/mol less with a 28-point difference in path lengths. For the POLARIS run, the option ‘Transition State Weighting’ was used with search depths {*r*_1_, *n*_7_}, {*r*_2_, *n*_5_}, and {*r*_4_, *n*_3_}; taking approximately 6 minutes using 4 processors.

For the ‘NODE BY NODE’ comparison, MEPSA’s node 2 (16, 23) and node 41 (47, 55) were selected from the set of MEPSA’s pre-defined nodes and chosen here based on their proximal similarity to the arbitrary points selected in the ‘GLOBAL’ comparison above. The pathway found by MEPSA returned an integrated energy of 28.5 kcal/mol with 98 points (Figure 6A). POLARIS used the same nodes 2 and 41 as user inputs and identified a 72-point pathway having an integrated energy of 20.0 kcal/mol (Figure 6B). The total difference between the two minimum energy approximations found by the two algorithms was 8.5 kcal/mol, again favoring POLARIS.

**Figure 6:**
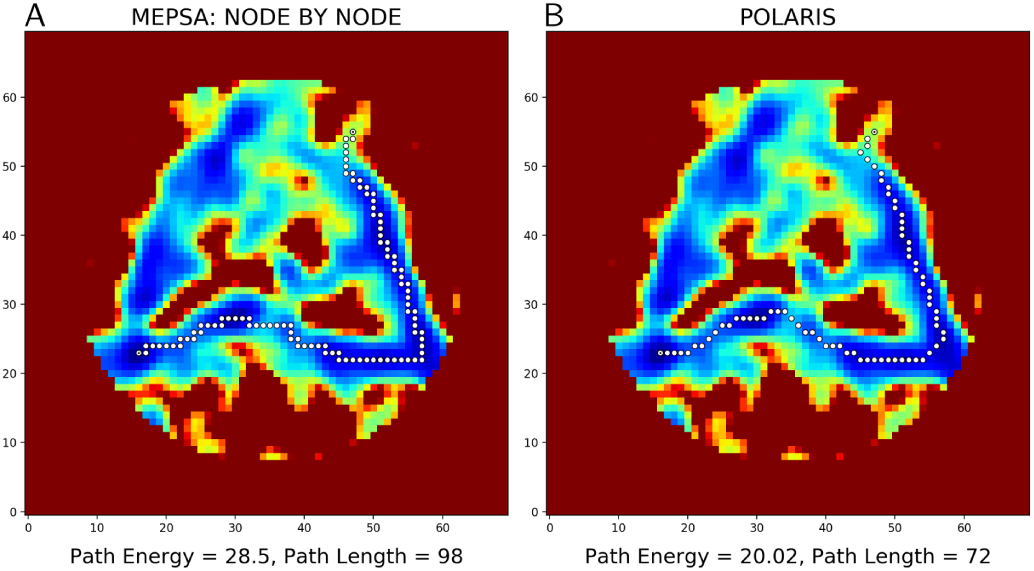
Comparison between MEPSA’s ‘NODE BY NODE’ algorithm (A) and POLARIS (B). While both algorithms again found the same approximate global path, POLARIS returned a path of integrated energy 8.5 kcal/mol less with a 26-point difference in path lengths. For the POLARIS run, the option ‘Transition State Weighting’ was used with search depths {*r*_1_, *n*_7_}, {*r*_2_, *n*_5_}, and {*r*_3_, *n*_3_}; taking approximately 3.5 minutes using 4 processors.

## DISCUSSION

In both comparisons with MEPSA, POLARIS’ outputs gave a substantially lower energy path while providing higher flexibility in the choice of user-defined start and end points, as well as in the number of optional user-defined transit locations. While the final paths from each follow the same broad regions throughout the landscape, the MEPSA algorithm appears to favor the production of jagged segments and deviates from POLARIS’ minimum path within regions of its locally defined nodes (such as in the southeast corner of the landscape).

As a further explanation for this large difference in energy and number of path points, MEPSA grows its trajectory in strictly cardinal directions (N, E, S, W), while POLARIS makes allowances for both cardinal and ordinal directivity (N, NE, E, SE, S, SW, W, NW) whenever locally required. Since the energy landscape ultimately represents a molecule’s movement as a combination of its two highest-ranking eigenvectors (the two reaction coordinates), any combination of the two must be accounted for - including cardinal (where only one eigenvector feature is altered at a time) and ordinal (where both features change simultaneously). In theory, this principle should hold for all reaction coordinates, regardless of context.

As for computation time, it should be noted that MEPSA generated the above paths within seconds. It should also be noted that its self-defined ‘global minimum’ solutions were undershot considerably by POLARIS’ techniques (which make no such claims to finding the ‘perfect’ solution). While POLARIS is capable of similar speeds at drastically lowered segmentation depth and permutational order, the lowest-energy trajectories found (shown here) required more computation time (10-20 minutes). Thus, when weighting accuracy over timing, POLARIS’ methods seem considerably more fit. Although these differences may seem trivial from a macroscopic view, such exactitude is essential on the microscopic level for accurately calculating biologically-relevant reaction rates via downstream algorithms. In the discussion to follow, POLARIS will be analyzed in terms of its ‘Completeness’, ‘Accuracy’, ‘Complexity’.

### Completeness

The greater the number of permutations that POLARIS is allowed to iterate through and compare against, the more accurate its computations will be in obtaining the minimum energy path. However, since the purpose of POLARIS is to limit this exhaustive search, it instead seeks to iteratively find the set of optimal approximations by use of feasible parameters (user-defined {^*N*^ *P*_*r*_} combinations) between each recursion, thus performing a series of computationally inexpensive operations that ultimately give rise to the same output as would one computationally exhaustive approach. Because of this, limits must be drawn on the total number of permutations allowed for the program to compute and compare between.

This total permutation limit is set by the user under the ‘Parameters’ section found within the ‘Settings’ tab (Figure 9). Here, combinations of *n* and *r* can be chosen for each 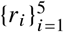, with the total number of permutations accessible from that combination calculated to the right (as given within the adjacent P(4^*n*^, r) entry box).

For *r*_1_, the maximum image subdivision depth, *n*_*max*_, is automatically chosen from the given datafile. At this depth, every value on the landscape is represented as a possible minimum node at least once, with this parameter thus comparing between all permutations of single points between each new *S* and *E* varying across all recursions. As *n*_*max*_ contains all other minima options within it (obtained individually from *n*_1_ to *n*_*max*−1_), it is redundant to compute any other combination less than this value for *r*_1_ (with equivalent logic holding for all other *r*_*i*_).

### Accuracy

Uncertainty is first introduced in the choice of the overhead metric and its ability to realistically encapsulate the dynamics of the system being explored. For example, for the potential energy surface of a chemical reaction, a reaction could be approximated either by the path having the lowest integrated energy, or by the path having the lowest maximum peak (activation energy). To provide flexibility in this overhead metric, POLARIS offers the ‘Transition State Weighting’ constraint, which can be enabled to weight the comparison of competing lowest-energy paths based on their rate-limiting step (point of maximal energy through which that path passes) instead of by just the net integrated energy along that path. When this feature is activated, POLARIS weights all energies across the landscape based on a power function - keeping lower energies approximately untouched while making higher-energy coordinates increasingly more unfavorable.

When bifurcation opportunities exist within a landscape (as in the case of the central, branching region of Figure 4), significant uncertainty can be introduced for the path of least action. This is especially relevant when such bifurcations are approximately degenerate - leading to two distinct paths of almost equivalent energy spanning radically different regions of the landscape.

During pathway comparisons on the example ribosome landscape, POLARIS isolated one such nearly equivalent energy bifurcation - dividing the ribosome’s most probable sequence of configurations into two discrete sets separated by a high-energy island (Figure 7). From an accuracy perspective, this instance illustrates the reliance on POLARIS’ user parameters in constructing its least-energy path, as well as the importance of the user experimenting with these parameters to achieve optimum results.

**Figure 7:**
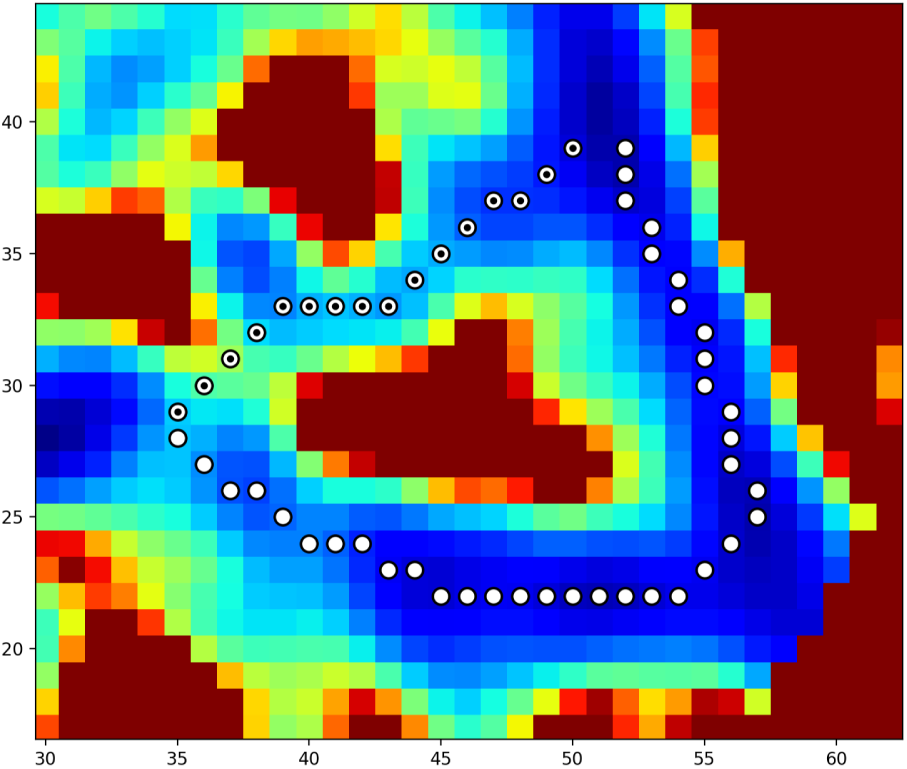
Comparison of two approximately degenerate paths found via POLARIS. To create each path, midpoints were placed alongside the initial user-defined start and end points - forcing POLARIS’ exploration of both routes independently on separate runs. These two diverging paths go through the center (black points) and southeast (white points) regions of the landscape and contain path energies 8.97 kcal/mol (with 16 points) and 8.50 kcal/mol (with 37 points), respectively. With only a difference of 0.47 kcal/mol between them, it is possible that such a degenerate least-energy bifurcation could represent novel conformational mechanics that allow for flexibility in macromolecular processes both spatially and temporally. For example, the bifurcation seen above may represent a shortcut in the ribosomal work cycle (elongation) that only becomes available under specific buffer or temperature conditions - allowing the ribosome to modulate its reaction rates based on fluctuating environmental signals.

As an aside, an exploration of such bifurcations can be instrumental for elucidation of a given system’s underlying dynamics. As such, the user can supply defined midpoints between the set of start and end coordinates to resolve such bifurcations (e.g., by placing one anchor at a minimum point in the middle of the landscape for one run, followed by another anchor instead at a minimum in the southeast corner for the next), with the added ability to then compare each least-energy approximation after completion of both runs.

### Complexity

In application, the number of contending lowest-energy paths become innumerable as the size and topological complexity of the landscape increases - making an exhaustive computational search for the path of least action infeasible towards these limits. Large and highly complex maps should therefore be used with this limitation in mind.

The range of available permutations to search through is proportional to the size of the data file (as seen via the available combinations of *n* and *r*). As increasingly larger combinations of {*n*_*i*_, *r* _*j*_} are chosen, the timing of each search will also increase, as defined by the number of available permutations (i.e., {^*N*^ *P*_*r*_}). Thus, the timing of each run is ultimately governed by the algorithm’s rate limiting step, `itertools.permutations`, via O(^*N*^ *P*_*r*_).

In its current state, POLARIS is best applied to energy landscapes of biologically-relevant size, at dimensions similar to the scale of reaction coordinates seen within chemistry and biophysics (19, 39). Here it has been shown that the algorithm is fully capable of representing the trajectories of highly complex structures within this domain (i.e., the ribosome, contained within a 70 × 70 dimension landscape). For landscapes larger than these recommended dimensions, pixel values could be binned beforehand or masked out to only those regions of interest - both of which will be supplied as future options within the POLARIS user interface.

## ACKNOWLEDGEMENTS

I would like to thank Suvrajit Maji and Hstau Liao for helpful comments on an earlier version of the manuscript. This research was supported by the National Institutes of Health Grants GM29169 and GM55440 (to J.F.).

## SUPPLEMENTARY MATERIAL

### User Interface

POLARIS supports Comma Separated Spreadsheet (.csv, .txt) inputs in the form of *m* × *n* heat maps. The GUI structure consists of a main window with two tabs entitled ‘Coordinates’ and ‘Settings’. Under the ‘Coordinates’ tab, an energy landscape can be imported and viewed. Once loaded, between 2 and 10 sets of user coordinates can be chosen, allowing the user to find the one-way path of least action between any two points, a series of points, or creating a cycle of points possibly corresponding to stable, reversible processes (Figure 8).

**Figure 8:**
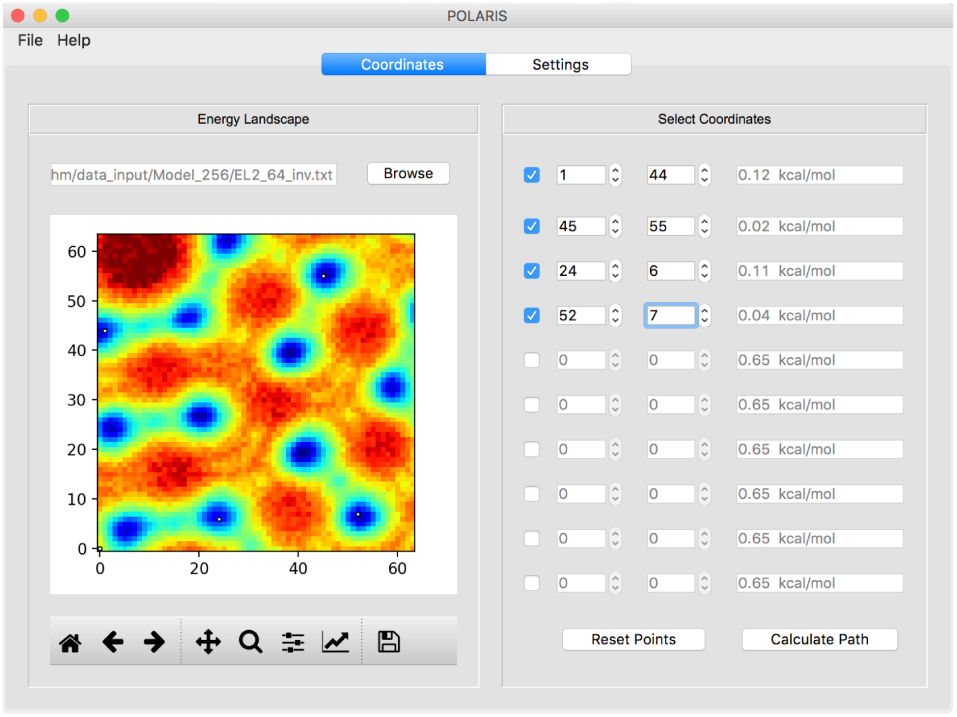
Image from the main page of the POLARIS user interface, allowing users to load a valid datafile and add up to 10 intermediate transit locations on the energy landscape as desired.

Advanced settings can be accessed via the ‘Settings’ tab, where parameters can be set to specify the algorithm’s maximum search depth and performance. The ‘Transition State Weighting’ option can also be enabled to weight POLARIS’ comparison of competing lowest-energy paths based their rate limiting step, as opposed to the integrated energy along all coordinates in that path. After these settings have been decided upon, the user can proceed back to the ‘Coordinates’ tab and click the ‘Calculate Path’ button to initiate the path finding algorithm.

Every value of *n* can be changed in the ‘Parameters’ section (Figure 9) by first unmarking its corresponding checkbox, if active. As that value of *n* is altered, the total permutation count will automatically update to the right. Adjusting these *n* parameters for each value of *r* such that the P(4^*n*^, r) values are all of the same order of magnitude will prevent any extreme rate limiting steps within the computation. Once these parameters have been set, they can be activated by checking each checkbox - thus instructing POLARIS to use those specific parameters within its search.

**Figure 9:**
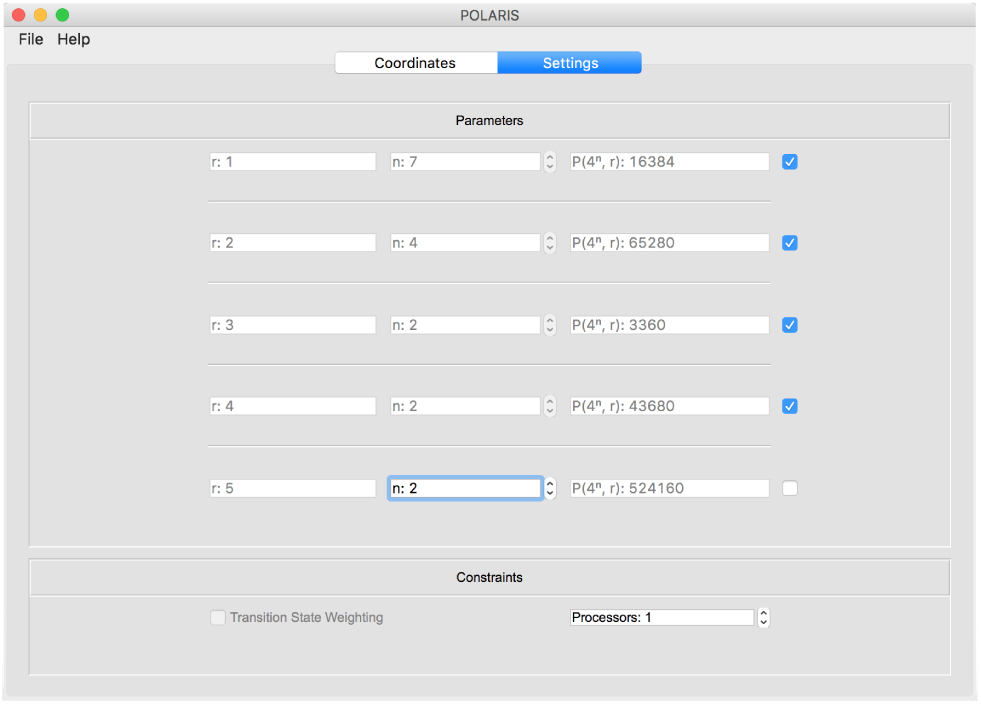
Image from the ‘Settings’ tab of the POLARIS user interface, allowing users to set parameters and constraints as desired. Note that each *n*_*j*_ has been chosen such that the total number of permutations for each {^*N*^ *P*_*r*_} combination are of approximately the same order of magnitude, and initially chosen as to be in line with the range provided by *n*_*max*_ (16,384 total permutations). As this order of magnitude is increased, so too will the computation time.

Upon completion of the backend algorithm, landscape-path plots (Figure 10) and transition state diagrams (Figure 11) are saved automatically as .png images. Additionally, coordinates of the minimum energy path and its respective energies are automatically generated within plain text files in the form of three column lists (*x, y, z* ∈ energy). These files also contain the total integrated energy, overall length of each trajectory, user defined parameters and elapsed computation time in their header.

**Figure 10:**
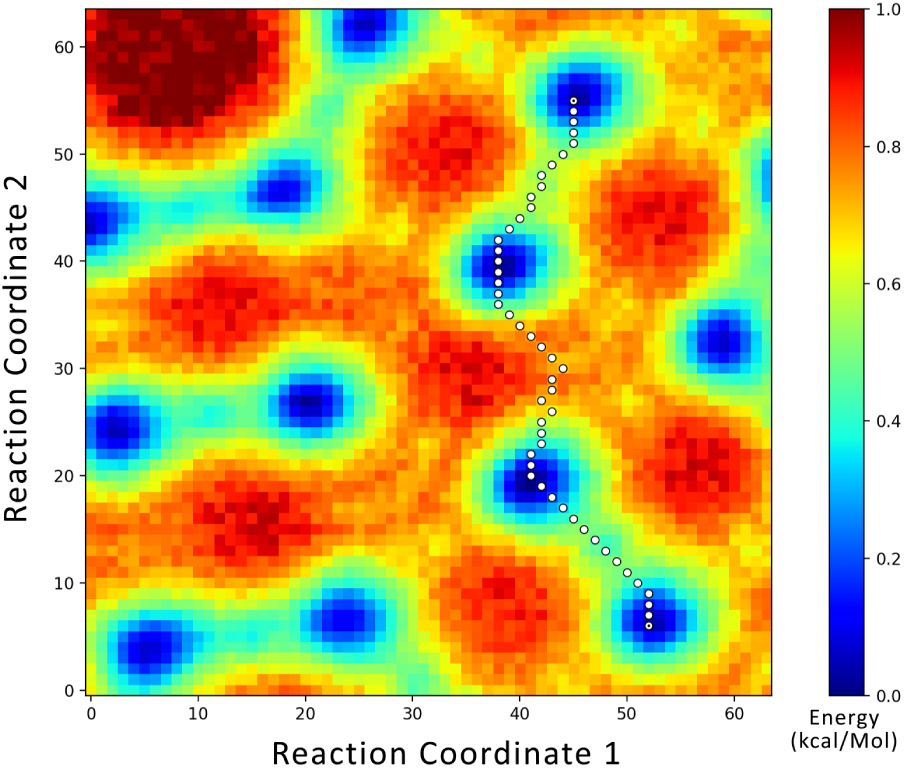
Example output of the least action path for an example, computationally-generated landscape.

**Figure 11:**
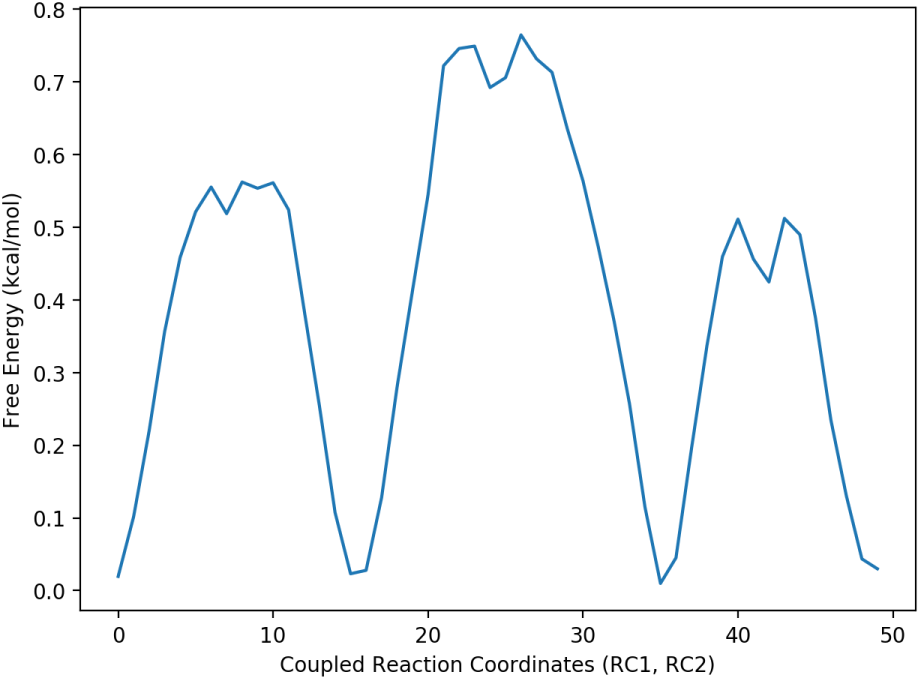
Example transition state diagram for the path seen in Figure 10, with energies plotted against the set of coupled coordinates (RC1, RC2).

## REFERENCES

1. Wales, D., 2003. Energy Landscapes. Cambridge University Press.

2. Errol, L., 2011. Computational Chemistry: Introduction to the Theory and Applications of Molecular and Quantum Mechanics. Springer, 2nd edition.

3. Whitford, P., R. Altman, P. Geggier, D. Terry, J. Munro, J. Onuchic, C. Spahn, K. Sanbonmatsu, and S. Blanchard, 2011. Dynamic views of ribosome function: Energy landscapes and ensembles. Springer, Vienna.

4. Munro, J., K. Sanbonmatsu, C. Spahn, and S. Blanchard, 2009. Navigating the ribosome’s metastable energy landscape. Trends in biochemical sciences 34 8:390–400.

5. Whitford, P., K. Sanbonmatsu, and J. Onuchic, 2012. Biomolecular Dynamics: Order-Disorder Transitions and Energy Landscapes, volume 75.

6. Weber, D., D. Bellinger, and B. Engels, 2016. New Algorithms for Global Optimization and Reaction Path Determination, volume 578.

7. Wales, D., and H. Scheraga, 1999. Global Optimization of Clusters, Crystals, and Biomolecules, volume 285.

8. Schramm, V., 1998. Enzymatic Transition States and Transition State Analog Design, volume 67.

9. Truhlar, D., B. Garrett, and S. Klippenstein, 1996. Current Status of Transition-State Theory, volume 100.

10. Pappu, R., R. Hart, and J. Ponder, 1998. Analysis and Application of Potential Energy Smoothing and Search Methods for Global Optimization, volume 102.

11. Frauenfelder, H., G. S. Sligar, and G. P. Wolynes, 1991. The energy landscapes and motions of proteins. Science 254:1598–1603.

12. Henzler-Wildman, K., and D. Kern, 2007. Dynamic personalities of proteins. Nature 450:964–972.

13. Henzler-Wildman, K., and D. Kern, 2009. Sending signals dynamically. Science 324:198–203.

14. Frank, J., 2006. Three-Dimensional Electron Microscopy of Macromolecular Assemblies: Visualization of Biological Molecules in Their Native State. Oxford University Press, Oxford, New York.

15. Sigworth, J. F., 2016. Principles of cryo-EM single-particle image processing. Microscopy 65:57–67. http://dx.doi.org/10.1093/jmicro/dfv370.

16. Frank, J., 2016. Generalized single-particle cryo-EM: a historical perspective. Microscopy 65:3–8. http://dx.doi.org/10.1093/jmicro/dfv358.

17. Frank, J., 2017. Nobel Lecture: Single-Particle Recon-struction - Story in a Sample..

18. Fischer, N., A. L. Konevega, W. Wintermeyer, M. V. Rodnina, and H. Stark, 2010. Ribosome dynamics and tRNA movement by time-resolved electron cryomicroscopy. Nature 466:329–333.

19. Dashti, A., P. Schwander, R. Langlois, R. Fung, W. Li, A. Hosseinizadeh, H. Y. Liao, J. Pallesen, G. Sharma, V. A. Stupina, A. E. Simon, J. D. Dinman, J. Frank, and A. Ourmazd, 2014. Trajectories of the ribosome as a Brownian nanomachine. Proceedings of the National Academy of Sciences 111:17492–17497. http://www.pnas.org/content/111/49/17492.

20. Chen, B., and J. Frank, 2016. Two promising future developments of cryo-EM: capturing short-lived states and mapping a continuum of states of a macromolecule. Microscopy 65:69–79. http://dx.doi.org/10.1093/jmicro/dfv344.

21. Frank, J., and A. Ourmazd, 2016. Continuous Changes in Structure Mapped by Manifold Embedding of Single-Particle Data in Cryo-EM 100:61–67. http://www.sciencedirect.com/science/article/pii/S1046202316300251.

22. Maupertuis, L. P., 1744. Accord de differentes loix de la nature qui avoient jusqu’ici paru incompatibles.

23. Euler, L., 1748. Reflexions sur quelques loix generales de la nature.

24. Dijkstra, W. E., 1959. Communication with an Automatic Computer http://www.cs.utexas.edu/users/EWD/PhDthesis/PhDthesis.pdf.

25. Cherkassky, V. B., V. A. Goldberg, and T. Radzik, 1996. Shortest paths algorithms: theory and experimental evaluation, volume Ser. A. 73 (2).

26. Hart, E. P., J. N. Nilsson, and B. Raphael, 1968. A Formal Basis for the Heuristic Determination of Minimum Cost Paths, volume SSC4. 4.

27. Floyd, R., 1962. Algorithm 97: Shortest Path, volume 5.

28. Cheung, G., E. Magli, Y. Tanaka, and M. K. Ng, 2018. Graph Spectral Image Processing. Proceedings of the IEEE PP.

29. Milanfar, P., 2013. A Tour of Modern Image Filtering: New Insights and Methods, Both Practical and Theoretical. IEEE Signal Processing Magazine 30:106–128.

30. Marcos-Alcalde, I., J. Setoain, J. I. Mendieta-Moreno, J. Mendieta, and P. Gómez-Puertas, 2015. MEPSA: minimum energy pathway analysis for energy landscapes. Bioinformatics 31:3853–3855. http://dx.doi.org/10.1093/bioinformatics/btv453. bioRxiv

31. Botea, A., M. Muller, and J. Schaeffer, 2004. Near optimal hierarchical path-finding. Journal of Game Development 1:7–28.

32. Rossum, G., 1995. Python Reference Manual. Technical report, Amsterdam, The Netherlands, The Netherlands.

33. PyQT, 2012. PyQt Reference Guide http://www.riverbankcomputing.com/static/Docs/PyQt4/html/index.html.

34. Hunter, D. J., 2007. Matplotlib: A 2D graphics environment. Computing In Science & Engineering 9:90–95.

35. Oliphant, E. T., 2015. Guide to NumPy. CreateSpace Independent Publishing Platform, USA, 2nd edition.

36. Kuzmin, P. Y., 1995. Bresenham’s Line Generation Algorithm with Built-in Clipping. Comput. Graph. Forum 14:275–280. http://dblp.uni-trier.de/db/journals/cgf/cgf14.html#Kuzmin95.

37. Spirin, A., 2009. The Ribosome as a Conveying Thermal Ratchet Machine, volume 284.

38. Whitford, P., S. Blanchard, J. Cate, and K. Sanbonmatsu, 2013. Connecting the Kinetics and Energy Landscape of tRNA Translocation on the Ribosome, volume 9.

39. Dashti, A., D. B. Hail, G. Mashayekhi, P. Schwander, A. d. Georges, J. Frank, and A. Ourmazd, 2018. Functional Pathways of Biomolecules Retrieved from Singleparticle Snapshots. bioRxiv https://www.biorxiv.org/content/early/2018/03/30/291922.

